# Lower Limb Vertical Stiffness and Frontal Plane Angular Impulse during Perturbation-Induced Single Limb Stance and Their Associations with Gait in Individuals Post-Stroke

**DOI:** 10.1101/2023.04.10.536288

**Authors:** Keng-Hung Shen, James Borrelli, Vicki L. Gray, Mark W. Rogers, Hao-Yuan Hsiao

## Abstract

**Background:** After stroke, deficits in paretic single limb stance (SLS) are commonly observed and affect walking performance. During SLS, the hip abductor musculature is critical in providing vertical support and regulating balance. Although disrupted paretic hip abduction torque production has been identified in individuals post-stroke, interpretation of previous results is limited due to the discrepancies in weight-bearing conditions.

**Objective:** To investigate whether deficits in hip abduction torque production, vertical body support, and balance regulation remain during SLS when controlling for weight-bearing using a perturbation-based assessment, and whether these measures are associated with gait performance.

**Methods:** We compared hip abduction torque, vertical stiffness, and frontal plane angular impulse between individuals post-stroke and healthy controls when SLS was induced by removing the support surface underneath one limb. We also tested for correlations between vertical stiffness and angular impulse during perturbation-induced SLS and gait parameters during overground walking.

**Results:** During the perturbation-induced SLS, lower hip abduction torque, less vertical stiffness, and increased frontal plane angular impulse were observed at the paretic limb compared to the non-paretic limb, while no differences were found between the paretic limb and healthy controls. Vertical stiffness during perturbation-induced SLS was positively correlated with single support duration during gait at the paretic limb and predicted self-selected and fast walking speeds in individuals post-stroke.

**Conclusions:** Reduced paretic hip abduction torque during SLS likely affects vertical support and balance control. Enhancing SLS hip abduction torque production could be an important rehabilitation target to improve walking function for individuals post-stroke.

## Introduction

Stroke is a leading cause of disability in the United States^1^. After a stroke, walking ability is a strong predictor of quality of life^2^. Restoring walking function is one of the most important goals in post-stroke rehabilitation^3^. Deficits in the paretic single limb stance (SLS), including shorter duration and reduced weight-bearing compared to the non-paretic side^4, 5^, are commonly observed during gait and limit walking performance post-stroke^6^. This asymmetric SLS profile is associated with a slower walking speed^4^ and a reduced balance capacity^7^. Thus, understanding the mechanism underlying impaired paretic SLS is an important step towards developing effective gait rehabilitation approaches.

During the single support phase in gait, the stance limb supports the bodyweight and contributes to controlling balance. Supporting the bodyweight requires adequate vertical stiffness regulation, typically measured as the ratio of the change in vertical ground reaction force (vGRF) with respect to the vertical displacement of the body center of mass (CoM)^8^. Without sufficient stiffness, limb collapse may occur with increased vGRF during single support and may increase the risk of falling^9^.

Thus, abnormal stiffness regulation could contribute to deficits in SLS post-stroke. Although sufficient vGRF is required to support the body, it also induces an external torque that rotates the body away from the stance limb. Normally, such a destabilizing torque is counteracted by the medially-directed ground reaction force (GRF) beneath the stance limb to control whole-body angular momentum (WBAM)^10, 11^. A reduced ability to regulate WBAM may indicate poor balance control^12, 13^ and lead to early termination of the single support phase or increased risk of falls during walking. Thus, both vertical stiffness and WBAM need to be considered during functional SLS.

Simulation studies have shown that the hip abductor muscles play an important role in providing bodyweight support^14^ and regulating WBAM^11^ during the single support phase in gait. Along with the hip abductors, extension torques generated from the ankle, knee, and hip joints are required to support the bodyweight during SLS^15, 16^. However, after stroke, voluntarily extending the paretic lower limb often simultaneously induces involuntary hip adduction at the paretic side, which is considered clinically to be an abnormal extensor synergy^17^. This synergistic torque production has also been quantitatively characterized in several studies^18–20^. In these studies, the participants were secured in postures that emulated the stance phase in gait and were instructed to produce extension or plantarflexion torque at the ankle, knee, or hip joints isometrically. Under such conditions, a reduced hip abduction torque production capacity was reported at the paretic limb compared to their non-paretic limb and to healthy controls. These findings provide a potential explanation for the impaired paretic SLS, as vertical support and balance regulation could be affected by this synergy-related diminished hip abduction torque production capacity. However, these studies used a device to fully support participants’ bodyweight and to restrain trunk motion. In addition, the magnitude of torque coupling was quantified relative to the maximum capacity of the tested limb^18–20^. Maintaining an upright standing posture without external body-weight support requires more complex multi-segmental coordination between lower limb joints and the trunk. More importantly, interpreting the torque production relative to the isolated maximum capacity may not fully capture the participants’ ability to perform functional tasks, as supporting a similar amount of bodyweight likely requires a different percentage of the maximum joint torque in the paretic versus non-paretic limb. These limitations likely hindered the ability to link the observed abnormalities during these relatively isolated assessments to functional gait deficits after stroke.

On the other hand, interpretation of abnormalities observed during gait after stroke, including reduced paretic hip abduction torque production^21^, altered lower limb joint stiffness^22^, and increased WBAM^12^, are oftentimes confounded by different amounts of weight-bearing between the lower limbs. This is because individuals post-stroke commonly have difficulty transferring bodyweight towards the paretic stance limb during gait^23, 24^, and hip abduction torque production, lower limb stiffness, and balance regulation demands likely vary with different weight-bearing magnitudes. Thus, an approach that studies SLS under more controlled weight-bearing conditions is needed to elucidate the potential factors affecting paretic SLS.

To address the aforementioned issues, we have developed a novel perturbation-based assessment that induces SLS during standing without external bodyweight support, while controlling for the weight-bearing symmetry. We investigated whether abnormal hip abduction torque production persists at the paretic limb during induced SLS in individuals post-stroke, and whether the paretic limb exhibits altered vertical stiffness and regulation of WBAM compared to the non-paretic limb and healthy controls. Additionally, we examined whether the vertical stiffness and WBAM measured during perturbation-induced SLS are associated with single support duration during walking and are predictive of walking speed in individuals post-stroke.

We hypothesized that, with symmetric initial weight-bearing between lower limbs, individuals post-stroke would show a reduced paretic hip abduction torque during the perturbation-induced SLS phase compared to the non-paretic side and healthy controls. We further hypothesized that individuals post-stroke would show a reduced vertical limb stiffness and greater change in WBAM (i.e. angular impulse) in the paretic limb than the non-paretic limb and controls during the perturbation-induced SLS. Lastly, we hypothesized that vertical stiffness and change in WBAM in the paretic and non-paretic limbs during perturbation-induced SLS will be associated with single support durations during walking and predict self-selected and fast walking speeds in individuals post-stroke.

## Method

### Participants

Fifteen individuals post-stroke (62.8±8.0yrs; 6 females; time post-stroke 13.5±12.7yrs; 3 right paretic) and fifteen age- and sex-matched healthy controls (64.2 ± 8.2yrs; 6 females) participated in this study. For the post-stroke group, the inclusion criteria were (1) had the most recent episode of stroke occurred more than six months prior to study enrollment, (2) able to stand without any support for five minutes, and (3) able to walk with or without a walking aid for at least ten meters, and the exclusion criteria were (1) advised against participating in regular exercise by a medical professional, (2) having underlying medical conditions that impacted their ability to walk beyond the effect of stroke, and (3) unable to follow instructions. Healthy controls were included if they had no self-reported musculoskeletal or neurological injuries. The study was approved by the University of Maryland Baltimore Institution Review Board and all participants provided written informed consent to participate.

### Experimental Protocol

Participants post-stroke were instructed to walk along an instrumented 7-meter GAITRite walkway (*CIR Systems Inc., US*) at their self-selected and fast walking speeds. The participants completed two walking trials for each speed. Participants post-stroke then completed the Step Test (ST) to assess mobility. Participants placed one foot onto a 7.5-cm-high step and then back to the floor repeatedly as fast as possible for 15 seconds^25^. Each participant completed 2 trials with each leg, and the number of steps completed in each trial was recorded. The highest ST score from each leg was reported in Table. 1.

**Table 1.**
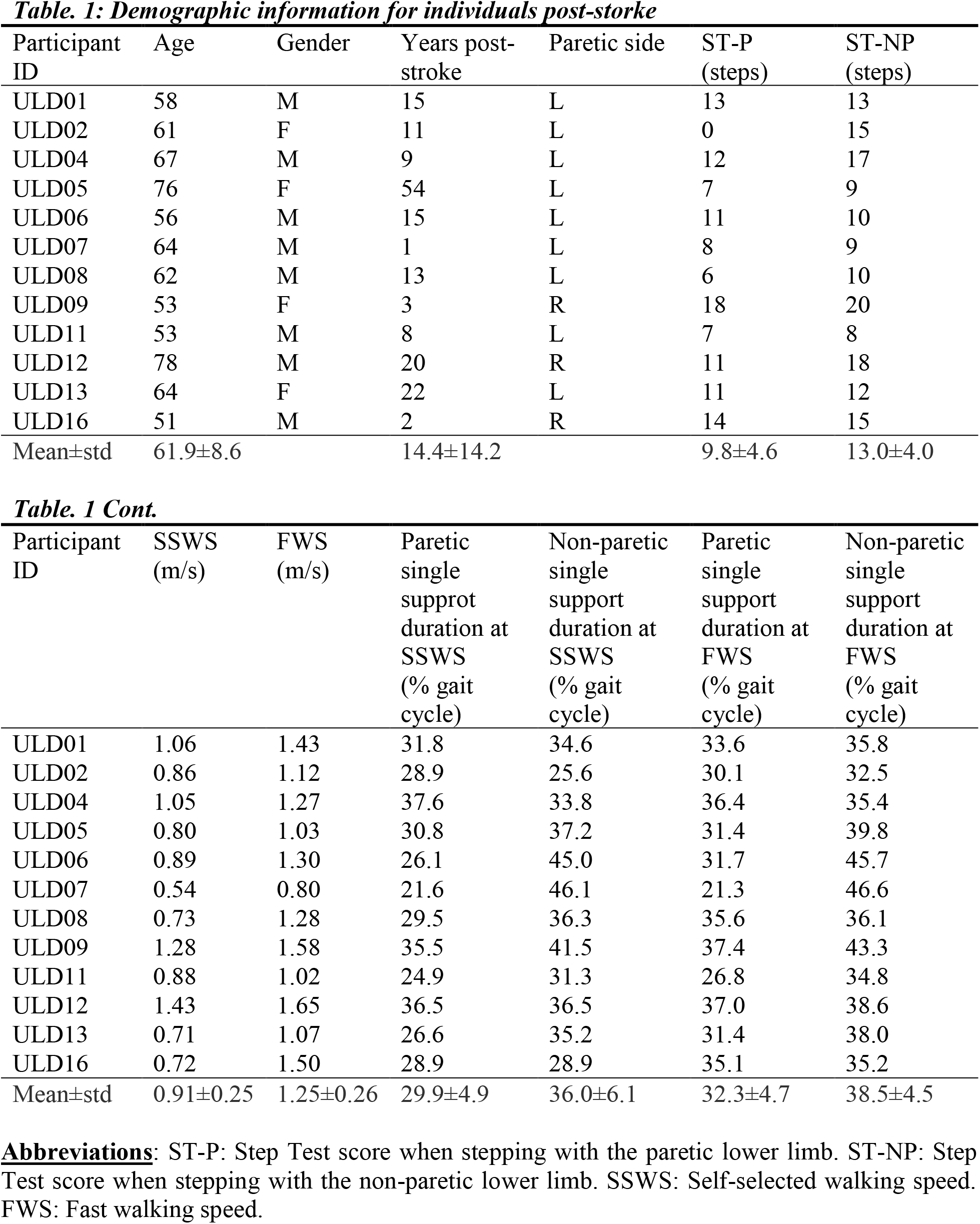
Demographic information for individuals post-storke.

### Perturbation-induced SLS

Two platforms with a height of 37 centimeters were each placed above a force platform (*Advanced Mechanical Technology Inc., US*) and adjacent to each other (*Fig.1*). The platforms consisted of a detachable surface where the participants could stand on, held by a frame using electromagnets (*Magnetech Corp., US*).

**Figure 1:**
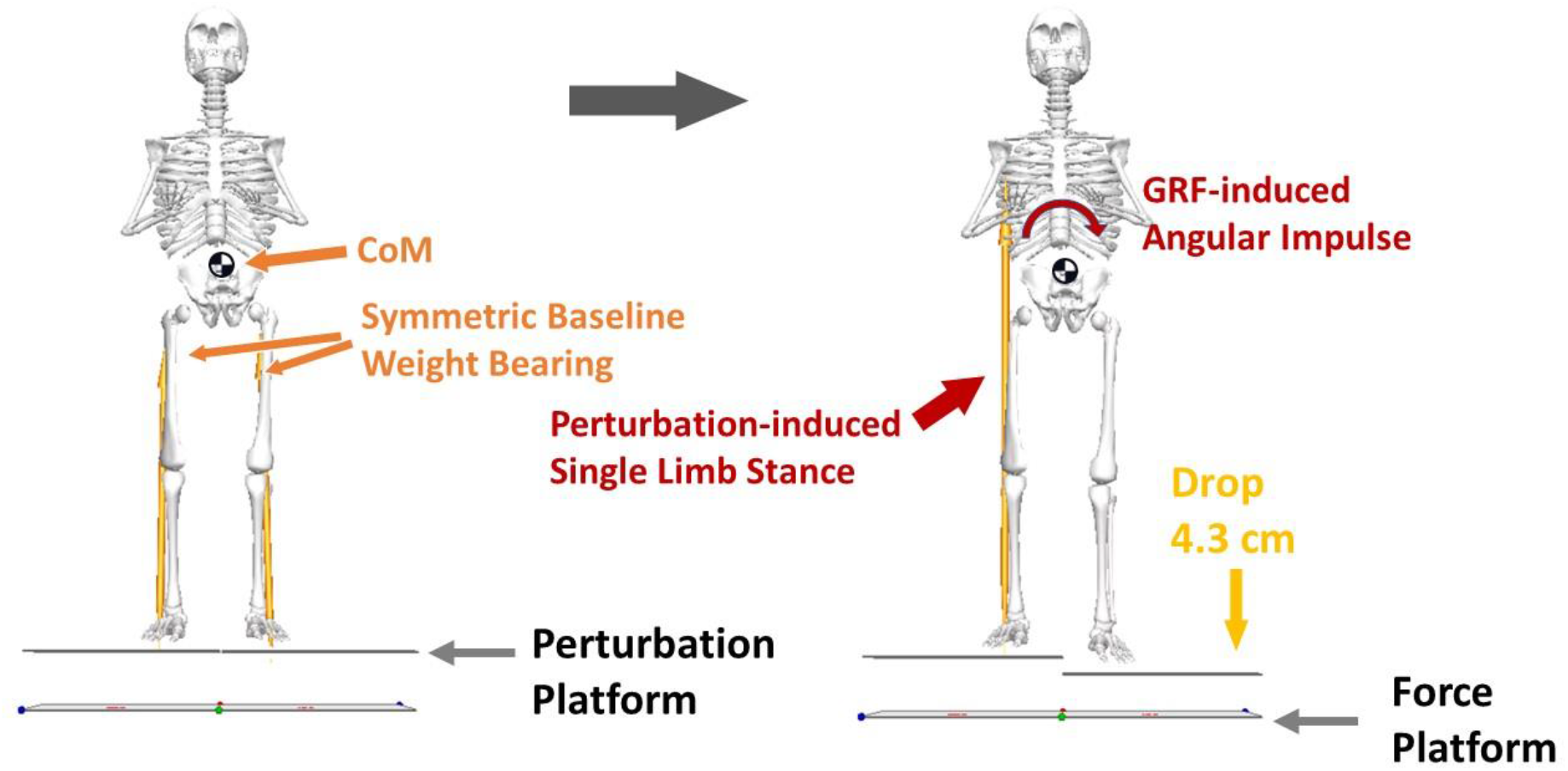
Schematic diagram of the perturbation-based assessment used to induce single limb stance (SLS).

Participants stood with one foot on each perturbation platform and were instructed to distribute their bodyweight evenly between legs. An experimenter monitored the real-time vGRFs to ensure even weight distribution before the perturbation was delivered. One of the two detachable surfaces was then released from the support frame upon disengagement of the magnets at an unpredictable time. The released support surface was dropped 4.3 centimeters vertically due to gravitational force. The non-dropped limb was induced into SLS during the aerial phase of the dropped support surface because minimum vertical support could be gained from the limb that was on the dropping platform.

The participants were instructed to react naturally to the perturbation and to maintain an upright posture. After one practice trial for each side, four perturbation trials were delivered to each side in a randomized order. The position of the foot was marked on the platform to ensure a consistent initial position throughout all trials.

All participants wore a gait belt during overgound gait assessment and ST, and an overhead safety harness during perturbation-induced SLS trials.

### Data Recording and Analysis

Overground walking speed and single support duration of the paretic and non-paretic limb were recorded and calculated using the GAITRite software during self-selected and fast walking speed trials.

During perturbation-induced SLS, a 10-camera motion capture system (*Oxford Metrics, UK*) was used to record the 3-dimensional position of 51 reflective markers placed on the participant and the perturbation platforms. Thirty-nine of the 51 markers were placed based on the Plug-in Gait marker set^26^. Ten additional markers were placed bilaterally on the fifth metatarsophalangeal joint, medial malleolus, femoral epicondyle, elbow, and greater trochanter. One marker was placed on each of the detachable surfaces of the perturbation platform. Two force platforms (*Advanced Mechanical Technology Inc., US*) located beneath the perturbation platform were used to record GRF and center-of-pressure (CoP) data. Kinematic and kinetic data were sampled at 150 Hz and 1500 Hz, respectively. Kinematic and kinetic data recorded during the perturbation-induced SLS trials were filtered offline at a cutoff frequency of 6 Hz and 30 Hz, respectively, with a zero-lag fourth-order Butterworth lowpass filter.

The SLS phase was defined as the aerial phase of the dropped platform. For a free-falling motion, the aerial phase could be estimated as (Eq.1):

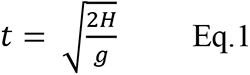

Where H is the drop distance (0.043 m), g is the gravity acceleration (9.81 m/s^2^), yielding a free-falling time of 0.094 s. Thus, the SLS phase was defined as the 94-millisecond window starting from perturbation onset. The aerial phase of the dropped platform was slightly longer than the calculated value for a freefall, likely due to friction between the dropped platform and its surrounding structure. The maximum displacement of the dropped platform at the end of the defined SLS phase was 0.041 m among all the trials, confirming that the 94-millisecond window did not capture any movement after ground contact of the dropped platform.

A 15-segment model (head, trunk, pelvis, and bilateral upper arm, forearm, hand, thigh, shank, and foot) was used to calculate the CoM position and frontal plane hip joint torque on the non-dropped side using Visual3D software (*C-Motion inc., US*). The GRFs and hip joint torque were normalized to bodyweight.

The change in WBAM was quantified as the angular impulse caused by the GRF about the COM in the frontal plane during the perturbation-induced SLS phase^27^. The angular impulse in the frontal plane was calculated by integrating the external moment with respect to time throughout the perturbation-induced SLS phase^27^ (*Fig.2C*). The external moment was defined as the cross product of the moment arm and the normalized GRF on the frontal plane, where the moment arm was defined as the vector from CoM to CoP and normalized to body height^11^. The direction of the external moment that rotates the body towards the dropped side was defined as positive.

**Figure 2:**
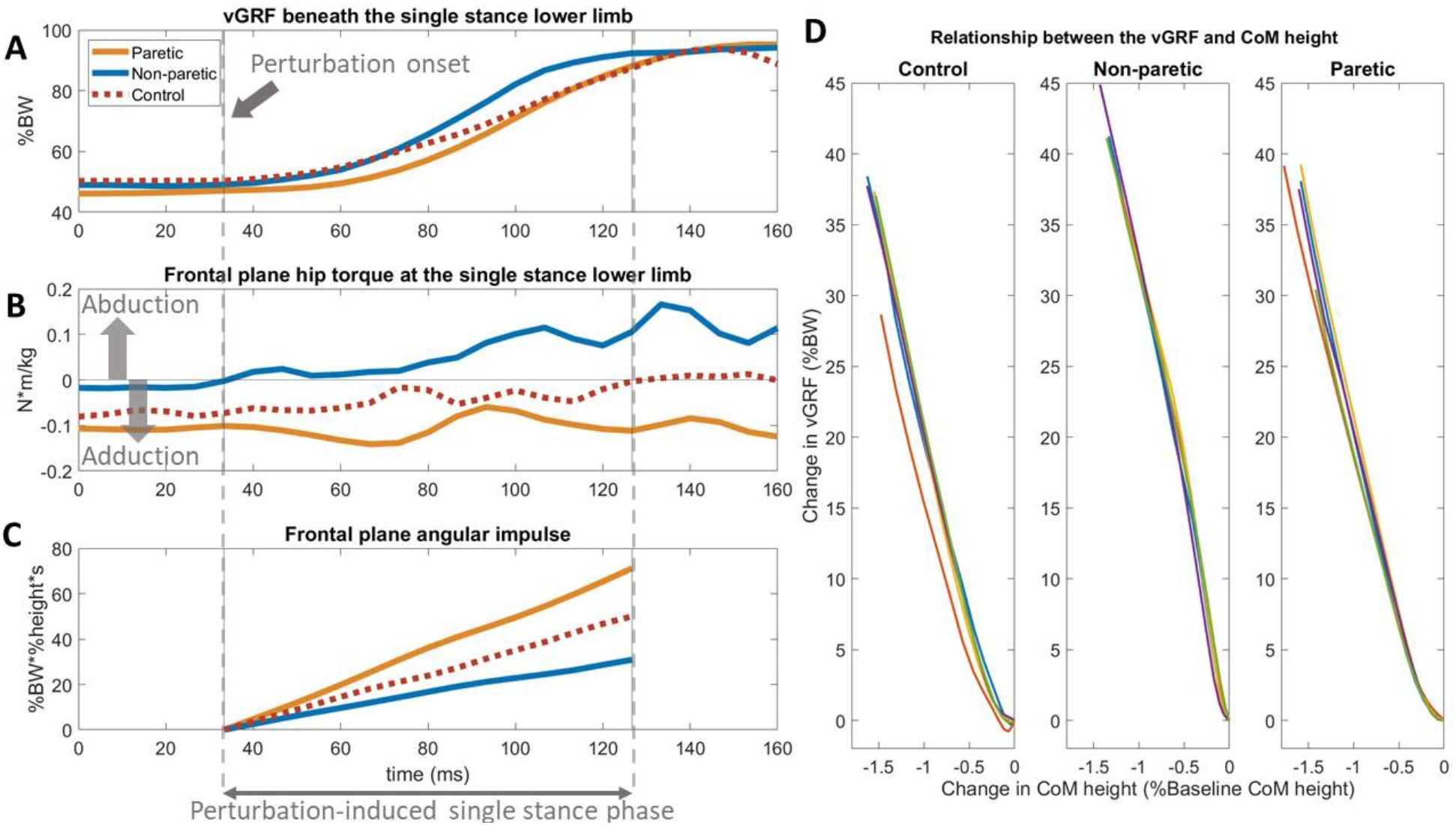
(A-C) A single trial of vGRF, hip torque, and angular impulse during perturbation-induced single limb stance phase in the paretic (orange solid line), non-paretic (blue solid line), and healthy control (red dotted line). (D) The change in vGRF and CoM vertical displacement during single limb stance phase from all trials of a representative participant in each group. Lines with different colors in the same subplot indicate different trials from the same single stance limb. vGRF: vertical ground reaction force. CoM: body center-of-mass.

Vertical stiffness was estimated as the slope of the regression line between the CoM vertical displacement and the change in the vGRF beneath the single stance (non-dropped) limb during the SLS phase^28^, where the CoM vertical displacement was normalized to the CoM height at perturbation onset. The coefficient of determination (*R*^2^) was used to quantify the appropriateness of fit^28^ (*Fig. 2D*). Vertical stiffness asymmetry (VSA) was calculated using (Eq.2).

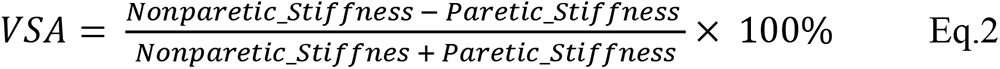

The baseline hip ab/adduction torque was defined as the average frontal plane hip joint torque of the single stance limb from 50 ms leading to the perturbation onset. The peak hip abduction torque of the single stance limb was identified during the SLS phase. The calculation of stiffness and angular impulse, and the extraction of baseline and peak hip abduction torque were performed in Matlab (*MathWorks, US*) with a customized algorithm.

### Statistical Analysis

Linear mixed effect models were used to test between-group differences in vertical stiffness, frontal plane angular impulse, and baseline and peak hip abduction torque during perturbation-induced SLS. For each dependent variable, we fitted a model that include a fixed effect of lower extremity (paretic, non-paretic, and non-dominant side in controls) and a random intercept of participant ID^29, 30^. The Wald F test was used to determine the significance of the fixed effect^31^, and Tukey’s HSD approach was used for post-hoc pairwise comparisons. Estimated marginal means and standard errors were presented in the results.

Pearson’s correlation was used to determine the associations between single support duration during overground walking and vertical stiffness and frontal plane angular impulse during perturbation-induced SLS in individuals post-stroke. Stepwise linear regression was performed to investigate whether vertical stiffness and frontal plane angular impulse during perturbation-induced SLS could predict self-selected and fast walking speeds. Data were converted into z-scores before stepwise model selection.

In the final regression model that predicts fast walking speed, vertical stiffness in paretic and non-paretic limbs were both significant predictors. This model implied that the variance of fast walking speed that was explained by vertical stiffness of individual limb could be explained by VSA because the beta coefficients for these two predictors had opposite signs. A nested model comparison was used to gain further insight into these results by comparing the two linear regression models that predicted fast walking speed, which the first model contained only VSA as the predictor, and the second model added vertical stiffness in paretic and non-paretic limbs as additional predictors to the first model. Data were converted into z-scores in both models.

All the statistical analysis was performed using R^32^. Alpha was set at 0.05 for all comparisons.

## Results

Ankle-foot-orthosis use may affect limb stiffness during perturbation-induced SLS, therefore the data from 2 participants who wore an ankle-foot-orthosis were excluded. In addition, data from 1 participant were excluded due to technical issues in the kinematics recording. After excluding those participants, data from 12 participants with chronic stroke (*Table. 1*) were analyzed.

### Perturbation-induced SLS

Prior to perturbation onset, all three groups produced adduction torque at the hip joint. The paretic limb showed a greater hip adduction torque compared to the non-paretic limb (*p<0.01; Fig.3A*).

**Figure 3:**
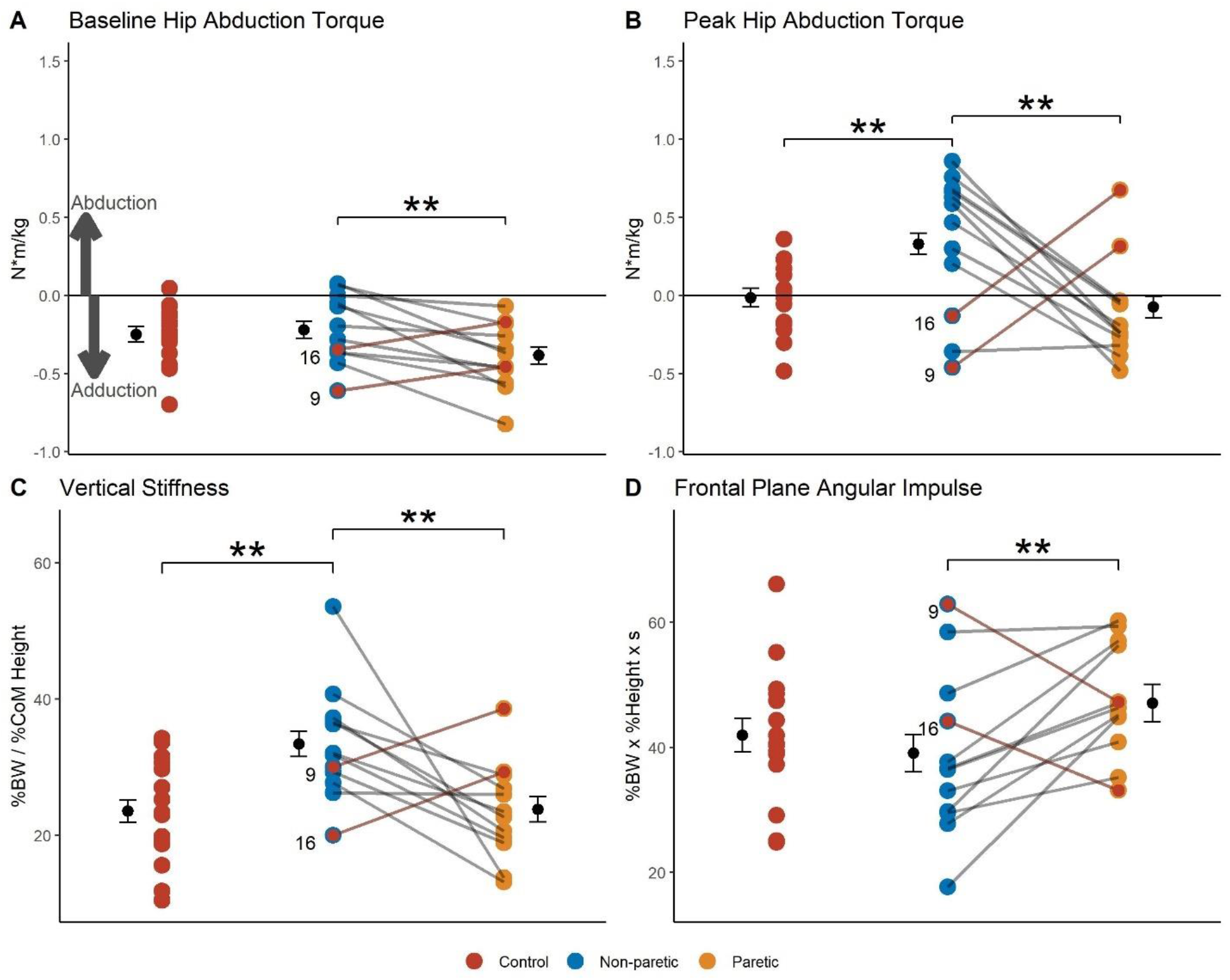
Between-group comparisons in perturbation-induced single limb stance assessment. Positive value indicates hip abduction in (A) and (B), and angular impulse that rotates the body away from the single stance limb in (D). Two participants (ULD09 and 16) that had an opposite trend in between-limb comparisons compared to other participants in individuals post-stroke are highlighted.

During the SLS phase, the non-paretic limb produced a larger peak hip abduction torque compared to the paretic limb and controls (*p-values<0.01; Fig.3B*).

When regressing the change in vGRF on the CoM vertical displacement during the SLS phase, the average *R*^2^ was over 0.97 for all three groups (controls: 0.97; paretic: 0.97; non-paretic: 0.98; *Fig.2D*). The vertical stiffness was greater in the non-paretic limb compared to the paretic limb and controls (*p-values<0.01; Fig.3C*).

The angular impulse in the frontal plane was greater in the paretic compared to the non-paretic limb (*p<0.01; Fig.3D)*.

Two participants post-stroke showed an opposite trend in between-limb comparisons compared to other participants. Specifically, they exhibited a greater hip abduction torque and vertical stiffness, and a lower angular impulse in the paretic versus non-paretic limb. These participants also had a greater fast walking speed among all individuals post-stroke (*Fig. 5D*) and are highlighted in Fig.3-5.

### Associations between perturbation-induced SLS and walking

In individuals post-stroke, vertical stiffness in the paretic limb during perturbation-induced SLS was positively correlated with single support duration when walking at self-selected (*r=0.59, p=0.04; Fig.4A)* and fast speeds (*r=0.66, p=0.02; Fig.4B*), while vertical stiffness in the non-paretic limb was not correlated with single support duration during either walking speeds (*Fig.4C-D*).

**Figure 4:**
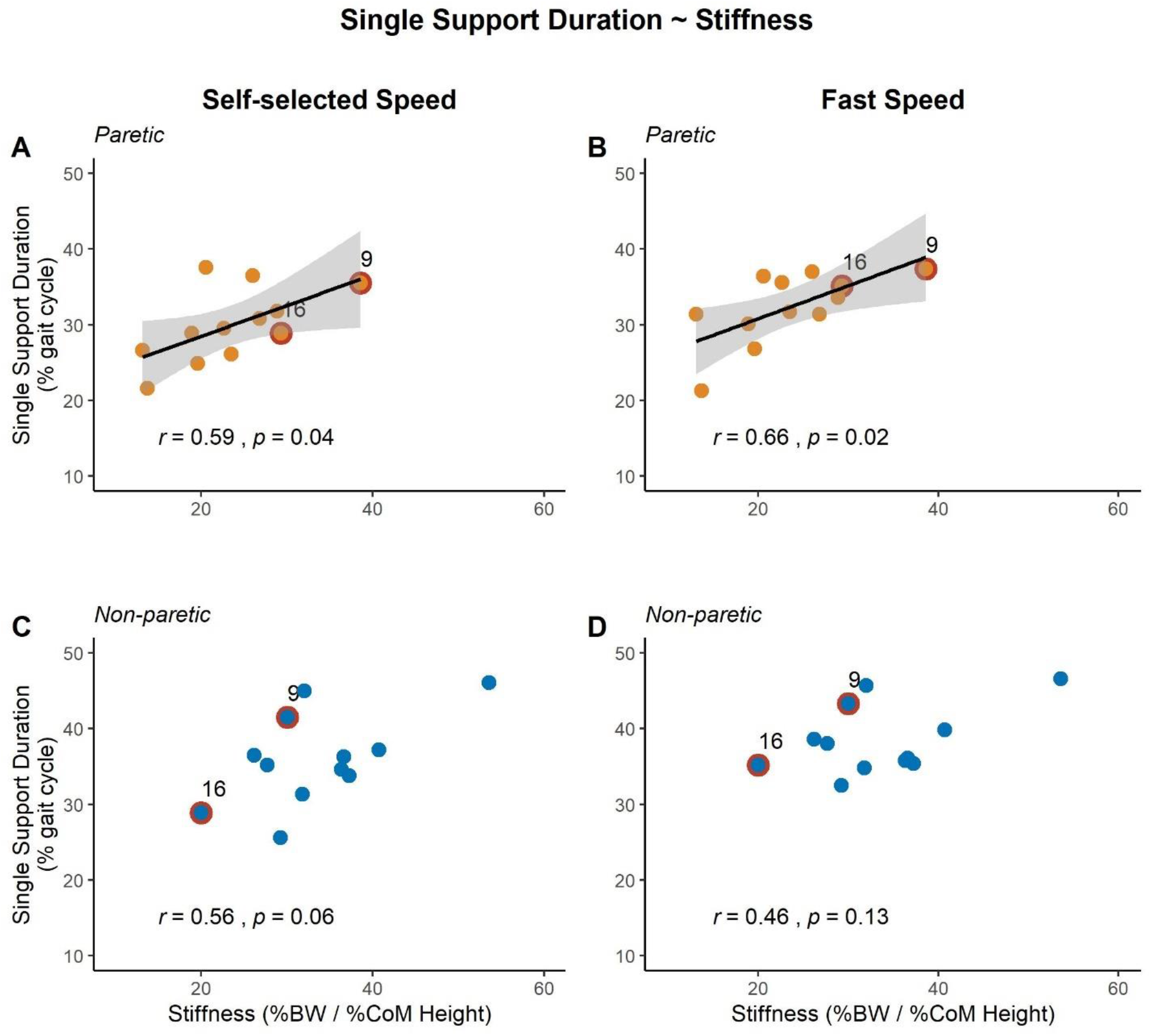
Correlations between vertical stiffness during perturbation-induced single limb stance phase (x-axis) and single support duration during overground walking (y-axis) in the paretic and non-paretic lower limb. The graphs on the top (A, B) and bottom (C, D) panels show the correlations in the paretic limb and non-paretic limb, respectively. The graphs on the left (A, C) and right (B, D) panels show the correlations with single support duration in self-selected and fast walking speeds, respectively. Two participants post-stroke (ULD09 and 16) that had an opposite trend in between-limb comparisons in the perturbation-induced single limb stance assessment compared to other participants are highlighted.

No correlation was detected between frontal plane angular impulse during perturbation-induced SLS and single support duration during either walking speeds (*p-values>0.19*).

### Predictors of walking speeds

When including vertical stiffness and angular impulse during both paretic and non-paretic SLS as predictors of walking speeds, vertical stiffness of the paretic limb was the only significant predictor and explained 36.3% variances of self-selected walking speed *(Fig.5A)*. Vertical stiffness of the paretic and non-paretic limb were both significant predictors that explained 74.6% variance of fast walking speed (*Fig.5B-D*).

**Figure 5:**
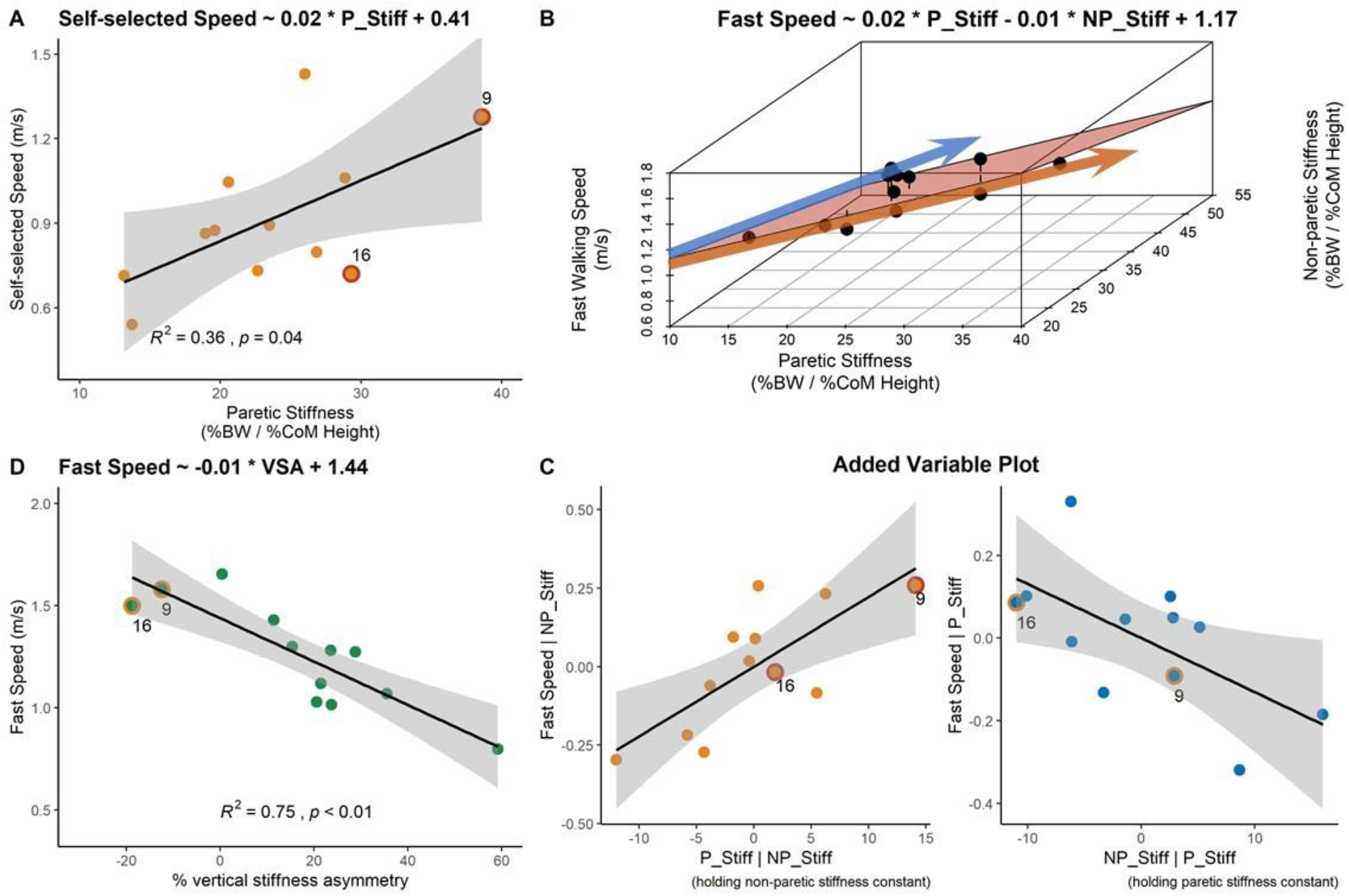
Prediction of overground walking speeds. Because only stiffness was included as predictors in the regression models, the beta coefficient from unstandardized data were shown to present the relationship in the variables’ original scales. (A) Vertical stiffness in the paretic lower limb predicted 36% of the variance in self-selected walking speed. (B) Vertical stiffness in the paretic and non-paretic lower limb together predicted fast walking speed, while the sign of beta coefficients had an opposite sign. (C) Added variable plot shows the unique contribution of variance in fast walking speed explained by vertical stiffness in each lower limb. (D) Vertical stiffness asymmetry predicted 75% of the variance in fast walking speed. Two participants post-stroke (ULD09 and 16) that had an opposite trend in between-limb comparisons in the perturbation-induced single limb stance assessment compared to other participants are highlighted.

The nested model comparison showed that compared to the model that contained only VSA as the predictor, adding vertical stiffness during paretic and non-paretic SLS did not significantly increase the variance explained in fast walking speed (*p=0.93*).

## Discussion

Using a novel support surface perturbation to induce SLS under controlled weight-bearing, we found that individuals post-stroke had reduced hip abduction torque, reduced vertical stiffness, and greater frontal plane angular impulse in the paretic limb compared to their non-paretic side. We also found that less vertical stiffness in the paretic limb during perturbation-induced SLS was associated with shorter paretic single support duration and slower self-selected walking speed. In addition, the asymmetry of vertical stiffness between lower limbs explained about 75% of the variance in fast walking speed.

### Hip Ab/Adduction Torque During Perturbation-induced SLS

Our results showed that individuals post-stroke exhibited greater adduction torque in the paretic hip joint compared to the non-paretic limb when standing upright with the bodyweight evenly distributed between the lower extremities. Since extension torques from lower limb joints are required for upright stance^15, 16^, the greater baseline adduction torque in the paretic limb observed in the present study compared to the non-paretic side could be explained by the synergistic extension-adduction torque coupling^18–20^.

During the perturbation-induced SLS phase, we observed a transition from hip adduction to abduction torque only for the non-paretic limb but not the paretic side. Similarly, in gait studies, a reduced paretic hip abduction torque during single support phase was observed in individuals post-stroke^21, 33^. However, interpretation of these observations was complicated by abnormalities in weight-bearing characteristics during walking. Specifically, individuals post-stroke oftentimes show limited weight transfer towards the paretic stance limb, therefore may require less hip abduction torque to support the bodyweight during the paretic stance phase compared to the non-paretic side^23, 24^. In the present study, weight-bearing conditions were controlled for between limbs. Thus, our findings advanced current knowledge by showing that between-limb asymmetry in hip abduction torque remain detectable in individuals post-stroke when weight-bearing demands are comparable across limbs.

### Vertical Stiffness During Perturbation-induced SLS

Compared to the non-paretic limb, we found that individuals post-stroke exhibited less stiffness in the paretic limb. Similarly, recent studies have shown less paretic ankle stiffness compared to non-paretic side during the stance phase in gait^22, 34^. However, our results were partially in contrast to the commonly observed increased resistance to passive movement, i.e., spasticity, in the paretic extremities after stroke^35, 36^. This could potentially be explained by the different weight-bearing conditions under which the stiffness was measured. Clinically, spasticity is often tested by applying a quick passive movement to the joint^37^. Under such conditions, a hyperactive short-latency stretch reflex could be a contributor to increased joint resistance. However, when performing functional movements that require weight-bearing by the lower limb, other deficits such as muscle weakness^38^ and the reduction in long-latency stretch reflex^39^ also contribute significantly to the regulation of joint stiffness, and the abnormal short-latency stretch reflex might thus affect joint resistance to a lesser degree^36, 40^. The hip abductor muscles function to counteract the gravitational force that accelerates the trunk and pelvis downward during SLS^41, 42^. Based on our findings, reduced hip abduction torque in the paretic limb likely contributed in part to the observed lower stiffness compared to the non-paretic limb during perturbation-induced SLS.

Furthermore, improvement in clinical spasticity measures often only results in minimal improvements in walking performance in individuals post-stroke. For example, significant improvement in the Ashworth Scale has been demonstrated as a result of managing ankle plantarflexor spasticity with botulinum toxin but not gait velocity^43^. Another ankle plantarflexor self-stretch program showed a greater improvement in the Tardieu Scale compared to conventional therapy, but the walking speed only improved by 0.04 m/s more than the conventional group^44^, which is not functionally meaningful^2^. This evidence suggested that the increased stiffness in the paretic limb measured under passive movement might not be the primary cause of the deficits observed during locomotion. Additionally, our results showed that the individuals who had less vertical stiffness during the perturbation-induced paretic SLS walked with shorter paretic single support duration. Thus, instead of increased passive joint stiffness due to spasticity, decreased lower limb stiffness during paretic SLS could be a key limiting factor of walking function after stroke.

### Angular Impulse During Perturbation-induced SLS

Our result showed that the GRF beneath the paretic limb induced a greater angular impulse about the body CoM in the frontal plane compared to the non-paretic limb during the perturbation-induced SLS phase. This is consistent with previous findings showing a higher average rate of change in WBAM (i.e., average external moment) during the paretic versus non-paretic SLS in walking adaptability tasks^12, 13^. The observed less paretic hip abduction torque may have contributed to the larger angular impulse during paretic versus non-paretic SLS because the hip abductor muscles serve an important role in regulating WBAM by counteracting the external moment induced by the vGRF^11^. Thinking at a segment level, reduced hip abduction torque during paretic SLS that is insufficient to resist the gravitational force may result in a net torque that rotates the trunk away from the stance limb^41^, leading to a higher trunk sway that deteriorates postural stability^45, 46^. This highlights the importance of training hip abductor torque production in rehabilitation focusing on vertical support in individuals post-stroke, as a disproportional improvement between the ability to provide vertical support and to regulate WBAM could potentially increase the risk of falling. Taken together, training hip abduction torque production under weight-bearing conditions could be an important rehabilitation target to improve paretic SLS function after stroke.

### Compare the Paretic Limb in Individuals Post-stroke and Controls During Perturbation-induced SLS Assessment

Contrary to our initial hypothesis, we did not find any differences in peak hip abduction torque, vertical stiffness, or angular impulse between the paretic limb in individuals post-stroke and healthy controls during perturbation-induced SLS phase. One explanation for this finding could be that the previously observed deficits in paretic SLS after stroke are dependent on controlled variables, including weight-bearing and weight transfer demands, in the present study. During gait, SLS happens after the bodyweight is transferred from the contralateral limb, and individuals post-stroke often show diminished weight transfer towards and less weight supported by the paretic lower limb^24^. However, it is not clear whether it is the deficit in performing weight transfer that leads to reduced weight-bearing at the paretic lower limb, or it is the inability to fully support the bodyweight during paretic SLS results in an unwillingness to transfer weight. Our perturbation approach eliminated the weight transfer demand and controlled for weight-bearing. We found that differences in SLS mechanics were not detectable between the paretic limb and healthy controls. These findings suggest that when performing SLS with a controlled weight-bearing demand, deficits in providing vertical support and regulating WBAM in the paretic limb were less pronounced when compared to the healthy controls. However, the between limb asymmetries were still detectable and associated with gait performance.

### Mechanics Identified During Perturbation-induced SLS Predict Walking Speed

Studies have suggested that modulation of leg stiffness is an important factor that contributes to stable gait^47–49^. Consistent with these studies, we found that the vertical stiffness during the perturbation-induced SLS explained variations in walking speed in individuals post-stroke. It is likely that the vertical stiffness revealed by our novel perturbation assessment in part reflected the ability to provide vertical support during gait, as people who had higher paretic vertical stiffness also had a longer paretic single support duration during gait.

We found that the vertical stiffness of the paretic and the non-paretic lower limb during the perturbation-induced SLS together explained 75% of the variances in fast walking speed. In this regression model, the sign of the beta coefficient for paretic stiffness was positive, while the sign was negative for non-paretic stiffness. We also found that during perturbation-induced SLS, the non-paretic limb in individuals post-stroke had a higher vertical stiffness compared to the healthy controls. Taken together, these findings may indicate that compensation pattern in the non-paretic lower limb could affect walking ability in addition to the deficits in the paretic limb. It has been shown that asymmetry in spatiotemporal gait metrics after stroke could represent compensatory mechanisms adopted during walking^50^. In particular, step length and stance time asymmetry after a stroke were associated with walking performance^51–53^. However, correcting those spatiotemporal asymmetries generally does not improve walking performance post-stroke^54, 55^, suggesting that spatiotemporal asymmetry in gait may be a consequence of other primary deficits. Simulation studies have suggested that lower limb stiffness could capture fundamental gait mechanics^47, 56^, and our results show that VSA was a strong predictor of fast walking speed among individuals post-stroke. These observations motivate future studies to investigate the relationship between changes in stiffness asymmetry and changes in walking speed following intervention.

### Limitations

There are limitations in the present study. First, the SLS duration induced by the perturbation is shorter than the single support duration during walking in individuals post-stroke^57^. This is a constraint in our approach as inducing a SLS duration that is typically observed during gait requires increasing the drop height in our perturbation paradigm. This would cause a large impact force to the dropped limb after the perturbation, which could be intolerable to the study participants. The short interval likely captured the responses that mostly arose from passive mechanisms, as any existing voluntary control would have minimal effect on mechanical behavior during the analyzed interval. Thus, our findings likely reflect changes in passive mechanisms that may have contributed to the abnormal gait pattern after stroke. Future development of novel experimental paradigms that induce a longer SLS interval may enable a more comprehensive investigation of deficits in paretic SLS post-stroke.

Additionally, the whole-body rotational dynamics (i.e., WBAM) analyzed in the present study might only reflect one aspect of balance regulation. Future study will investigate the interaction between WBAM and other biomechanical balance measures that could reflect whole-body translational dynamics^58^, and how different segments, including pelvis and trunk, contributed to the change in WBAM, to provide a more comprehensive picture of balance regulation during SLS.

## Conclusion

Using a perturbation approach to induce SLS while controlling for weight-bearing, we demonstrated that individuals post-stroke generated hip adduction torque during paretic SLS, while the non-paretic stance limb produced hip abduction torque. This abnormal paretic hip torque production was accompanied by reduced vertical stiffness and higher angular impulse in the paretic stance limb compared to the non-paretic side. In addition, vertical stiffness during perturbation-induced SLS phase was associated with single support duration during gait and walking speeds. However, no differences in perturbation-induced SLS mechanics were detected between the paretic limb in individuals post-stroke and healthy controls. Our findings suggest that under similar weight-bearing conditions, deficits in paretic SLS, including reduced hip abduction torque, altered stiffness, and greater WBAM, became less pronounced compared to healthy controls than generally thought, while performance asymmetry between the limbs was still present in individuals post-stroke and was associated with walking function. Improving hip abduction torque generation during paretic SLS may be an important target for gait rehabilitation post-stroke.

## Acknowledgement

The authors gratefully acknowledge the assistance of Robert Creath in development of the perturbation device and the experimental protocol, and the assistance of Doug Pizac, Breanna Arbuco, and Nathan Frakes in data collection and processing.

